# TPPB modulates PKC activity to attenuate neuroinflammation and ameliorate experimental multiple sclerosis

**DOI:** 10.1101/2024.02.02.578637

**Authors:** Shruthi Shanmukha, Wesley H. Godfrey, Payam Gharibani, Judy J. Lee, Yu Guo, Xiaojing Deng, Paul A. Wender, Michael D. Kornberg, Paul M. Kim

## Abstract

Protein kinase C (PKC) plays a key role in modulating the activities of the innate immune cells of the central nervous system (CNS). A delicate balance between pro-inflammatory and regenerative activities by microglia and CNS-associated macrophages is necessary for the proper functioning of the CNS. Thus, a maladaptive activation of these CNS innate immune cells results in neurodegeneration and demyelination associated with various neurologic disorders, such as multiple sclerosis (MS) and Alzheimer’s disease. Prior studies have demonstrated that modulation of PKC activity by bryostatin-1 (bryo-1) and its analogs (bryologs) attenuates the pro-inflammatory processes by microglia/CNS macrophages and alleviates the neurologic symptoms in experimental autoimmune encephalomyelitis (EAE), an MS animal model. Here, we demonstrate that (2S,5S)-(E,E)-8-(5-(4-(trifluoromethyl)phenyl)-2,4-pentadienoylamino)benzolactam (TPPB), a structurally distinct PKC modulator, has a similar effect to bryo-1 on CNS innate immune cells both in vitro and in vivo, attenuating neuroinflammation and resulting in CNS regeneration and repair. This study identifies a new structural class of PKC modulators, which can therapeutically target CNS innate immunity as a strategy to treat neuroinflammatory and neurodegenerative disorders.

## Introduction

Aberrant neuroinflammation by the innate immune cells of the central nervous system (CNS) contributes significantly to demyelination and neurodegeneration in multiple sclerosis (MS) and other neurologic disorders, e.g., Alzheimer’s disease (AD), Parkinson’s disease, and amyotrophic lateral sclerosis (Kutzelnigg and Lassmann, 2014; Hickman *et al*., 2018; Reich, Lucchinetti and Calabresi, 2018; Xie *et al*., 2022). In progressive MS, where the adaptive immune response plays a less prominent role, microglia and CNS-associated macrophages are activated in a pro-inflammatory phenotype that promotes demyelination and neurodegeneration (Mahad, Trapp and Lassmann, 2015; Voet, Prinz and van Loo, 2018; Faissner *et al*., 2019). Current Food and Drug Administration (FDA)-approved MS drugs primarily target peripheral lymphocytes and thus are effective in treating relapsing-remitting MS. However, these drugs inadequately address the disease progression and treatment of progressive MS by failing to modulate CNS innate immune cells. Therefore, compounds that therapeutically target CNS innate immunity to promote remyelination and CNS repair are greatly needed.

Protein kinase C (PKC) is a family of important signaling molecules that are involved in numerous cellular activities (Platten *et al*., 2003; Noh *et al*., 2012; Chen *et al*., 2014; Newton, 2018). Studies have demonstrated that the PKC isoforms in innate immune cells are important in key functions for stimulating CNS regeneration and repair, such as phagocytosis, secretion of neurotrophic factors, and providing supportive milieu (Zheleznyak and Brown, 1992; Hortelano, Genaro and Boscá, 1993; Smith *et al*., 1998; Fronhofer, Lennartz and Loegering, 2006; Kooij *et al*., 2015; Lim, Sutton and Rao, 2015). PKC is also a key signaling molecule in the triggering receptors expressed on myeloid cells 2 (TREM2) pathway, which is involved in critical microglia activities (Ulland and Colonna, 2018; Andreone *et al*., 2020; Kim and Kornberg, 2022; Wang *et al*., 2022). Thus, deletion or inhibition of this signaling pathway results in harmful outcomes. People with TREM2 missense mutation, Nasu-Hakola disease, experience an early onset of dementia and severe demyelination (Tanaka, 2000; Dardiotis *et al*., 2017).

We recently demonstrated that modulation of PKC activity hinders pro-inflammatory responses from microglia and macrophages, while activating anti-inflammatory and regenerative responses from these cells (Kornberg *et al*., 2018; Abramson *et al*., 2021; Gharibani *et al*., 2023). In experimental autoimmune encephalomyelitis (EAE), an MS animal model, modulation of PKC activity prevented the development of neurologic symptoms, and therapeutic treatment of EAE mice with a PKC modulator significantly improved the symptoms of EAE. Surprisingly, even in very late-stage EAE, when the peripheral inflammatory process has mostly subsided and only CNS inflammation persists, PKC modulation continued to ameliorate EAE symptoms. We also demonstrated using a lysolecithin (LPC)-induced demyelination model that modulation of PKC activity in microglia and CNS macrophages promoted remyelination by activating a regenerative phenotype in these cells, which provided a favorable anti-inflammatory environment, enhanced phagocytosis, and released beneficial factors to stimulate oligodendrocyte (OL) differentiation.

Our previous studies have focused primarily on modulating PKC activity with bryostatin-1 (bryo-1) and bryostatin analogs, termed bryologs (Abramson *et al*., 2021). Bryo-1, a natural compound that is CNS-penetrant and is a potent PKC modulator (Wender *et al*., 1988; Sun and Alkon, 2006; Hongpaisan, Sun and Alkon, 2011), was originally tested as a cancer drug because it inhibited the tumorigenic properties of phorbol esters, which activate PKC (Clamp and Jayson, 2002), and since then, bryo-1 has also been tested for other indications, including AD, immunotherapy augmentation, and HIV eradication (Bullen *et al*., 2014; Laird *et al*., 2015; Gutiérrez *et al*., 2016; Farlow *et al*., 2019; Hardman *et al*., 2020; Sloane *et al*., 2020). We established that bryologs have similar biological effects as natural bryo-1 on innate immune cells. We also demonstrated that bryologs alleviated the symptoms of EAE and that the activity of bryo-1 and its analogs require PKC binding. Furthermore, our initial study showed that prostratin and prostratin analogs also had similar effects in cultured macrophages (Abramson *et al*., 2021), indicating that other PKC modulators with chemical structures distinctly different from bryo-1 and bryologs may have therapeutic potential in treating neuroinflammatory and neurodegenerative disorders.

Here, we explored further the therapeutic potential of a structurally novel PKC modulator in stimulating an anti-inflammatory/regenerative phenotype in innate immune cells and in enhancing remyelination. We have identified (2S,5S)-(E,E)-8-(5-(4-(trifluoromethyl)phenyl)-2,4-pentadienoylamino)benzolactam (TPPB) (Kozikowski *et al*., 1997), a benzolactam that, while structurally different from bryo-1 and prostratin, binds to the C1 domain of PKC and has similar anti-inflammatory effects as bryo-1 and prostratin in cultured microglia and macrophages. We also demonstrate that TPPB, but not prostratin, improved EAE symptoms like bryo-1 and enhanced remyelination in a focal demyelination model. Our findings demonstrate that 1) TPPB has similar in vitro and in vivo effects to bryo-1 and 2) in vitro screening techniques allow for rapid screening of potential PKC modulators, but in vivo models, e.g., EAE and/or LPC-induced demyelination models, are required to establish the clinical potential of these compounds.

## Materials and Methods

### Mice

Wild-type C57BL/6J mice were purchased from the Jackson Laboratory (stock #000664). Animals were housed in a Johns Hopkins animal facility and acclimatized in the facility for at least one week. All animal experimental protocols were approved by the Johns Hopkins Institutional Animal Care and Use Committee.

### Induction and Scoring of EAE

EAE was induced in 8-12-week-old C57BL/6J female mice by subcutaneous immunization of myelin oligodendrocyte glycoprotein 35-55 peptide (MOG_35–55_). Briefly, 100 µg MOG_35–55_ was emulsified with 100 µl complete Freund’s adjuvant, and 50 µl of the emulsion was subcutaneously injected into each of two sites on the lateral abdomen on day 0. Additionally, on day 0 and day 2, mice received 250 ng of pertussis toxin intraperitoneally (IP). Clinical signs of EAE were assessed daily beginning on day 7 post-immunization. Scoring was performed in a blinded manner according to the following scale: 0, no clinical deficit; 0.5, partial loss of tail tone; 1.0, complete tail paralysis or both partial loss of tail tone plus awkward gait; 1.5, complete tail paralysis and awkward gait; 2.0, tail paralysis with hind limb weakness evidenced by foot dropping between bars of cage lid while walking; 2.5, hind limb paralysis with little to no weight-bearing on hind limbs (dragging), but with some movement possible in legs; 3.0, complete hind limb paralysis with no movement in lower limbs; 3.5, hind limb paralysis with some weakness in forelimbs; 4.0, complete tetraplegia but with some movement of head; 4.5, moribund; and 5.0, dead.

### Treatment of Mice with TPPB and SUW014

TPPB (Tocris, 5343) and SUW014 (provided by P.A.W.) were dissolved in DMSO to 1 mM stock solutions, which were further diluted to a desired concentration in sterile PBS for animal experiments. Mice were treated three days per week by IP injection of TPPB (50 nmol/kg), SUW014 (35 nmol/kg), or an equal volume of vehicle control. Mice were randomized and received their first treatment on the day they reached a clinical score of 1.0 or greater; thereafter, a three-days/week treatment schedule was followed.

### Isolation and Treatment of Murine Bone Marrow-Derived Macrophages (BMDM)

Briefly, 8-10-week-old C57BL/6J mice were euthanized, and bone marrow cells were isolated from their femurs in an aseptic environment by flushing with sterile PBS. The cell pellet was then collected by centrifugation (1500 rpm for 8 min). The red blood cells (RBC) were lysed in RBC lysis buffer, followed by centrifugation (1500 rpm for 8 min). The resulting cell pellet was resuspended in RPMI media supplemented with 10% FBS, 1% penicillin/streptomycin, 2 mM L-glutamine, 50 µM 2-mercaptoethanol, and 20 ng/ml recombinant mouse GM-CSF (Peprotech). The cells were incubated at 37°C for seven days to induce macrophage differentiation. Subsequently, on day 8, BMDM were replated and treated as needed for the experiments. For treatment of cultured cells, TPPB was dissolved in DMSO to a final concentration of 1 mM, which was subsequently dissolved in PBS or cell culture media for the required treatment. An equal volume of DMSO diluted in PBS or cell culture media was used as vehicle control.

### Preparation of Tissue for Flow Cytometry

Mice were euthanized with an overdose of isoflurane and then perfused with ice-cold PBS delivered via a cardiac puncture. Spinal cords were mechanically dissociated, then chemically dissociated with collagenase (200 U/ml) and DNase (100 U/ml) with constant shaking for 10 min, triturated with a pipette, and incubated for an additional 10 min. Cells were then passed through a 100-μm filter and washed with PBS. Myelin debris was removed by resuspending the cell pellet with a debris removal solution (Miltenyi Biotec), overlaying with PBS, and spinning at 3,000 g, and then removing the myelin debris layer. Cell pellets were resuspended in PBS, passed through a 100-μm filter, and stained with antibody as described below. Myeloid cell panels proceeded immediately to flow cytometry staining after cell isolation. For panels involving T cell cytokines, cells were first resuspended in a solution of complete RPMI with brefeldin/monensin and PMA/ionomycin for 4 hrs at 37°C.

### Flow Cytometry Staining

All staining was performed in the dark at room temperature. Cells were first stained with zombie NIR (1:1500) for 10 min along with CD16/32 (1:100) dissolved in PBS. Cells were then washed and resuspended in a solution of PBS+2% FBS+1 mM EDTA and stained with the relevant antibodies. All surface antibodies were stained at a 1:300 dilution. Cells were then washed and incubated in FoxP3 fixation/permeabilization buffer following the manufacturer recommended protocol. Intracellular staining was performed with conjugated antibodies (1:200) against the specified proteins in permeabilization buffer for 1 hr, washed twice, and then analyzed cells with a Cytek Aurora Flow cytometer. Data analysis was performed with FlowJo software.

### Enzyme-linked Immunosorbent Assay (ELISA)

BMDM were treated overnight with LPS (100 ng/ml) with or without TPPB (100 nM). Culture supernatants were collected, and cytokine production was assessed using ELISA kits for IL-12 (#88-7121-22), and IL-10 (#88-7105-22) purchased from eBioscience and following the manufacturer’s instructions. Plates were read at 450 nm on a plate reader.

### Western Blot

BMDM were treated overnight with LPS (100 ng/ml) ± TPPB (100 nM) or IL-4 (20 ng/ml) ± TPPB (100 nM). Cells were lysed in RIPA buffer supplemented with protease inhibitors, and protein concentration was determined by BCA assay. Protein samples were then prepared in SDS sample buffer and resolved by SDS/polyacrylamide gel electrophoresis. The bands were transferred to PVDF Immobilon-P membranes (Millipore) and blocked in TBS-T containing 5% BSA for 1 hr at room temperature. The membranes were probed overnight at 4°C with rabbit monoclonal antibodies against inducible nitric oxide synthase (iNOS; Cell Signaling Technology, 2982) or arginase-1 (Arg-1; Cell Signaling Technology, 93668), followed by incubation with horseradish peroxidase (HRP) conjugated with anti-rabbit secondary antibodies (Cell Signaling Technology, 7074). Protein was detected with SuperSignal chemiluminescent substrate solution (Pierce), and Western blots were visualized using the LI-COR imaging system. The protein loading of each sample was verified by stripping the membrane with Restore Western blot stripping buffer, blocked again as described above, probed for 1 hr at room temperature with HRP-conjugated anti-actin antibody (GeneScript, A00730), and visualized the protein expression as described earlier. The Western blot was quantified using ImageJ software.

### In Vitro Phagocytosis Assay with E. coli Bioparticles in BMDM

BMDM were treated overnight with LPS (100 ng/ml) with or without TPPB (100 nM). This was followed by aspirating the culture medium, and 100 µl of pHrodo Green E. coli BioParticles (ThermoFisher Scientific, P35366l) suspended in culture media were added to BMDM at a concentration of 50 bioparticles per cell. The cells were incubated for 3 hrs at 37°C in a humidified atmosphere of 5% CO_2_, washed twice with sterile PBS, and then incubated with 100 µl of diluted trypan blue with PBS (1:2 dilution) to quench the extracellular fluorescence. The cells were washed twice with PBS fixed with 100 µl of 4% paraformaldehyde (PFA), incubated for 15 min at room temperature, and washed with PBS (100 µl per well for each wash). Following the wash step, 50 µl of PBS was added to each well, and fluorescence intensity was measured on a SpectraMax fluorescent plate reader.

### Myelin Isolation and In-cell Western Blot (ICW) for Myelin Phagocytosis in BMDM

#### Myelin Isolation

Myelin isolation and labeling with carboxyfluorescein succinimidyl ester (CFSE) were performed as previously described (Rolfe *et al*., 2017), and 8-10-week-old C57BL/6J mice were used to isolate myelin. Briefly, the whole brains were dissected from euthanized animals and homogenized using a sterile hand-held rotary homogenizer in ice-cold 0.32 M sucrose solution. The homogenate was carefully overlaid onto ice-cold 0.83 M sucrose solution in a 50-ml polypropylene centrifuge tube, forming a sucrose density gradient, and centrifuged at 100,000 g at 4°C for 45 min in a pre-cooled ultracentrifuge rotor. The resulting white crude myelin interface was collected from the interface between two sucrose densities and further homogenized in a sterile hand-held rotary homogenizer for 30-60 sec before subjecting to centrifugation at 100,000 g at 4°C for 45 min in a pre-cooled ultracentrifuge rotor. The solid white myelin pellet was resuspended in Tris-Cl solution and centrifuged at 100,000 g at 4°C for 45 min. Subsequently, the pellet was resuspended in sterile PBS and centrifuged at 22,000 g for 10 min at 4°C. The final isolated myelin pellet was resuspended in sterile PBS to a final concentration of 100 mg/ml.

#### Myelin Labeling

For CFSE labeling, 10 mg of myelin was resuspended in 200 µl of 50 µM CFSE and incubated at room temperature for 30 min in the dark, followed by centrifugation at 14,800 g for 10 min at 4°C. The pellet containing labeled myelin was washed in 100 mM glycine in PBS for three times and resuspended in sterile PBS.

#### ICW for Myelin Phagocytosis in BMDM

BMDM were cultured on a clear bottom 96-well cell culture plate at a density of 2.5×10^4^ cells per well. The cells were treated with either vehicle or TPPB (100 nM) for 3 hrs, followed by incubation with 1 mg/ml of CFSE-labeled myelin for overnight at 37°C. BMDM were washed several times with sterile PBS to remove non-engulfed myelin debris. The cells were fixed with 4% PFA for 15 min and incubated with 0.1% Triton X-100 in PBS for 10 min. The fixed cells were incubated for 2 hrs at room temperature with β-actin antibody (Cell Signaling Technology, 4967) at 1:1000 dilution and then incubated with VRDye 549 Goat anti-Rabbit IgG Secondary Antibody (LI-COR biosciences, 926-54020) for 30 min. The plates were scanned on the LI-COR Odyssey imaging system according to the manufacturer’s protocol. The total fluorescence of ICW for CFSE-myelin was quantified and normalized to β-actin on ImageJ.

### Focal Spinal Cord Demyelination

#### LPC-induced Demyelination

Focal demyelination was induced in 8-12-week-old C57BL/6J mice by injecting 1% LPC (Sigma) in PBS into the ventral funiculus as described previously (Gharibani *et al*., 2023). Briefly, mice were anesthetized with ketamine (100 mg/kg) and xylazine (10 mg/kg) injected IP. Mice were placed on a stereotaxic frame, and a small midline incision was made below the ears on the back of the animal in the caudal direction. The prominent T2 was used as a landmark to expose the spinal cord and identify T3-4 intervertebral space. The dura was removed with a 32G needle without damaging the tissues. LPC (0.5 µl) was microinjected stereotaxically, at a rate of 0.25 μl/min using a microinjection syringe pump (UMP3; World Precision Instrument), into the right ventral funiculus (depth of 1.3 mm) via a 34G needle (Hamilton Co.) connected to a 10 µl Hamilton syringe. To avoid LPC efflux, there was a 2-min pause after LPC injection before retracting the needle. A single absorbable suture (Vicryl 5-0) was used to close the muscle and adipose tissues, following which the skin incision was closed with rodent wound clips. Saline (1 ml) and buprenorphine SR (1mg/kg; ZooPharm, LLC) were administered subcutaneously after the surgery to prevent dehydration and pain, respectively. Gentamycin (2 mg/kg; Henry Schein Animal Health) was given subcutaneously every 12 hrs for 3 days. Treatment with TPPB (50 nmol/kg, three times a week) or vehicle started 48 hrs after the surgery by IP injection.

#### Spinal Cord Harvest

The animals were anesthetized and transcardially perfused with PBS briefly, followed by ice-cold 4% PFA (Sigma). Spinal cord tissues were then harvested and made into a 3-mm piece of lesion site with the epicenter (site of injection) in the middle. The tissues were kept in 4% PFA at 4°C overnight for post-fixation, followed by cryoprotection in gradient sucrose (10% then 30% sucrose in PBS at 4°C overnight). The tissues were then embedded in O.C.T. and were coronally sectioned into 12-µm thickness sections at 0.8 mm from the epicenter at both sides (rostrally and caudally) using a cryostat (Thermo Shandon). The sections were collected on Superfrost Plus slides (VWR International) and stored at −80°C until staining.

#### Immunofluorescence Staining

The spinal cord sections were retrieved by Universal Antigen Retrieval Kit (R&D Systems) in a steamer for 15 min at 100°C, permeabilized with 1% Triton X-100 in TBS for 5 min, and incubated in blocking solution (10% donkey serum, 0.25% Triton X-100 in TBS) for 1 hr at room temperature. Primary antibodies were diluted in blocking solution and incubated for two overnights at 4°C. In order to study remyelination, double staining was performed with rabbit anti-Olig2 (1:100, Millipore) and mouse anti-APC (CC1) (1:100, Sigma). This was followed by incubating with appropriate fluorochrome-conjugated secondary antibodies (1:500, Thermo Fisher Scientific) for 2 hrs at room temperature, counterstaining with DAPI (3 µM in PBS; Sigma), and covering with antifade mounting media before placing coverslips. Minimum of six sections (3 rostral and 3 caudal) were counted manually by investigators who were blinded to the study groups. The total numbers of positive cells were normalized to the area of the injury using ImageJ.

## Results

### TPPB Induces an Anti-inflammatory Phenotype in BMDM

We have previously demonstrated that the PKC modulator bryo-1 promotes anti-inflammatory/regenerative activation of macrophages while concurrently suppressing pro-inflammatory markers in myeloid cells (Kornberg *et al*., 2018; Abramson *et al*., 2021). We also showed that the structurally unrelated PKC modulator SUW014, an analog of prostratin, exhibits bryo-1-like effects on myeloid cells, suggesting that these and related PKC modulators may serve as candidate drugs for targeting the innate immune cells in CNS. Here, we aimed to determine the anti-inflammatory potential of TPPB (Figure 1A), a cell-permeable high-affinity PKC modulator, on peripheral BMDM. BMDM were stimulated with LPS (100 ng/ml) for 24 hrs with or without treatment with TPPB (100 nM), and cytokine secretion in the cell culture supernatant was examined by ELISA. Treatment with TPPB significantly inhibited the production of the pro-inflammatory cytokine IL-12 induced by LPS (Figure 1B) and increased the secretion of the anti-inflammatory cytokine IL-10 (Figure 1C). Furthermore, TPPB inhibited the expression of iNOS, a key marker of pro-inflammatory phenotype, induced by LPS and augmented the expression of Arg-1, a marker of anti-inflammatory/regenerative phenotype, after IL-4 stimulation (Figure 1D). Overall, these data suggest that TPPB promotes the activation of macrophages that are involved in the regulation of the immune responses to limit inflammation and promote tissue repair mechanisms.

**Figure 1:**
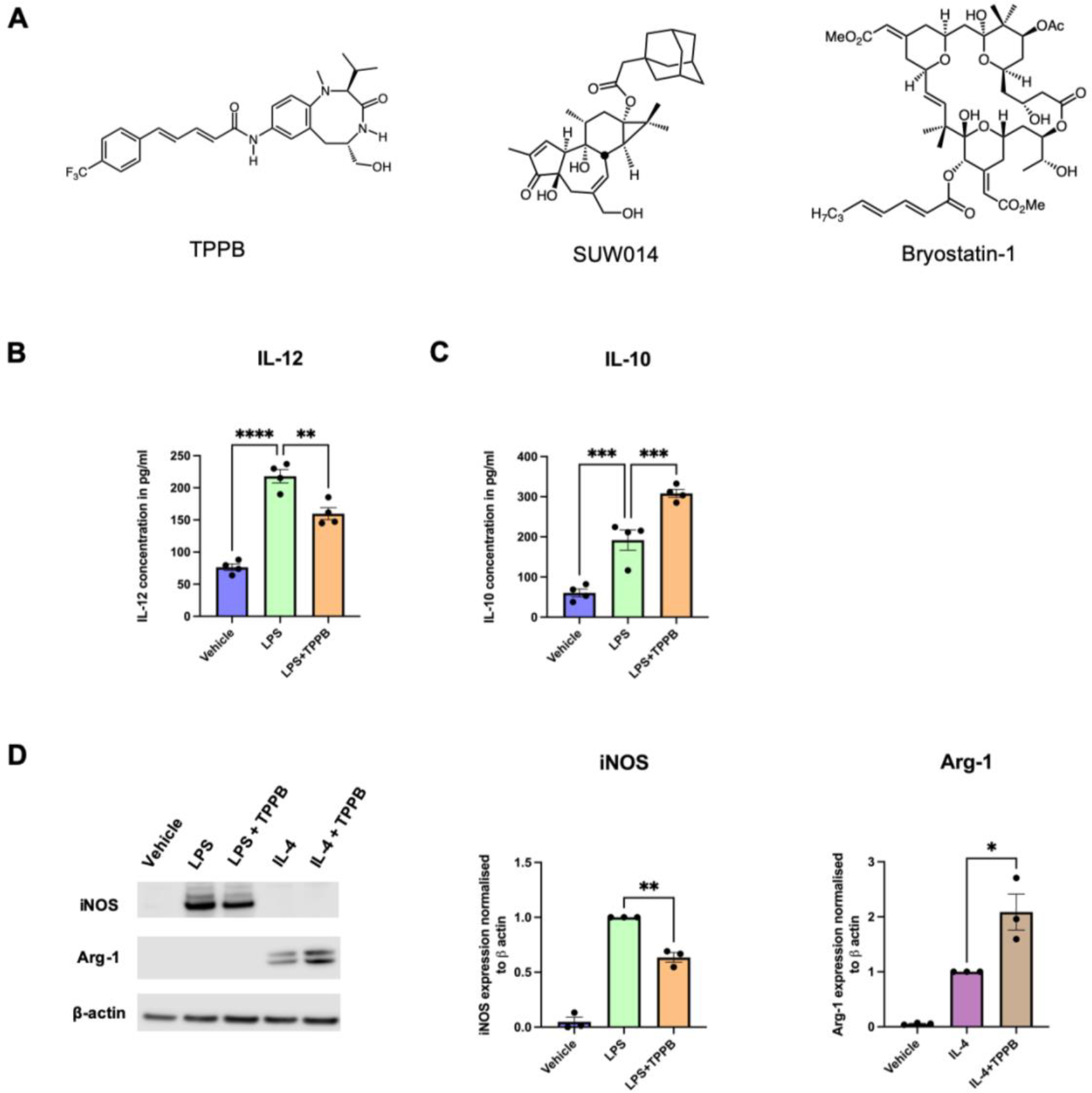
TPPB promotes an anti-inflammatory phenotype in BMDM. (A) Chemical structures for TPPB, SUW014 (prostratin analog), and bryo-1. BMDM were treated overnight as indicated in the figure with LPS (100 ng/ml), IL-4 (20 ng/ml), and TPPB (100 nM). Secretion of IL-12 (B) and IL-10 (C) into the culture media was measured via ELISA; n=4 experiments performed in replicates of three. (D) Representative Western blots of iNOS and Arg-1 (*left*) along with the histograms showing their quantification normalized to β-actin; n=3 for iNOS and Arg-1. The statistical differences between groups were calculated using one-way ANOVA for multiple-group analyses. All error bars depict SEM; p-value of <0.05 was considered significant (* p<0.05; ** p<0.01; *** p<0.001; and **** p<0.0001).

### TPPB Attenuates Neurologic Deficits in EAE

Our previous study demonstrated that bryo-1 attenuated the development and progression of EAE, and thus, bryo-1 represented a potential therapeutic agent for MS (Kornberg *et al*., 2018; Abramson *et al*., 2021). Similarly, having evaluated the anti-inflammatory effect of TPPB in vitro, we used the EAE model to carry out in vivo investigations of TBBP to better understand its clinical potential. MOG_35-55_-induced EAE model was used, and drug treatment began when the animals demonstrated EAE symptoms (clinical score of at least 1). In this experiment, we also tested prostratin analog SUW014, which was not done in the previous study (Abramson *et al*., 2021). We discovered that SUW014 (35 nmol/kg) had no effect on attenuating EAE symptoms when administered IP three days a week, and both vehicle- and SUW014-treated mice progressively developed neurologic symptoms (Figure 2A). On the other hand, TPPB (50 nmol/kg) attenuated the clinical symptoms of EAE and showed lower peak clinical score when administered IP three days per week at the onset of symptoms (Figure 2B). These results suggest that TPPB, like bryo-1, provides a beneficial effect on EAE mice and serves as a promising candidate for MS treatment.

**Figure 2.**
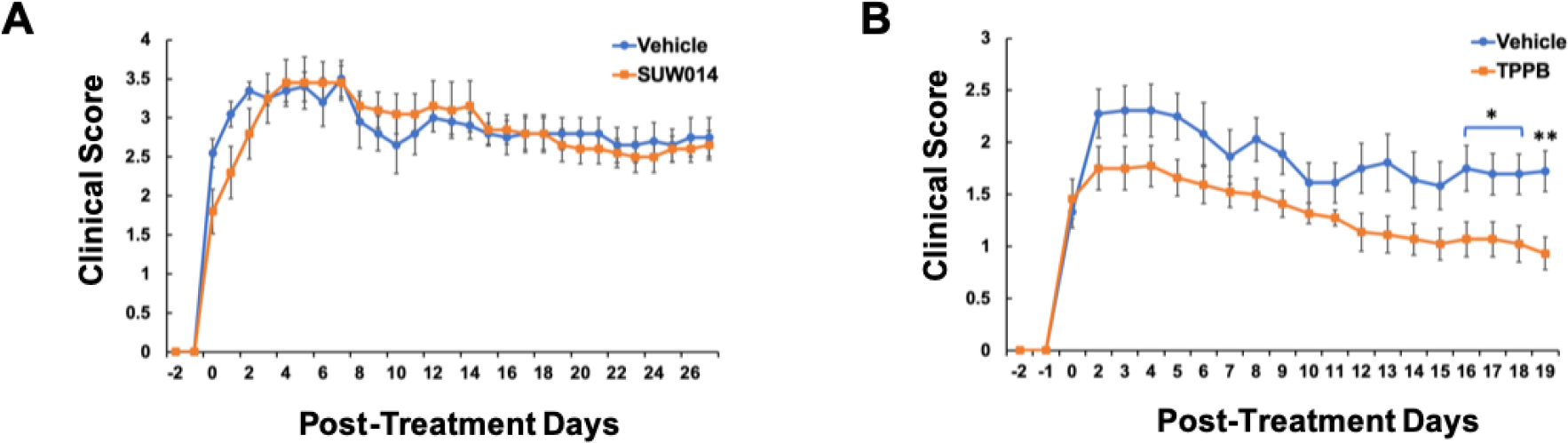
TPPB attenuates neurologic deficits in MOG_35-55_-induced EAE mice. Mice were treated with SUW014 (A) and TPPB (B) by IP injection three times a week at the onset of EAE symptoms. TBBP significantly ameliorated the neurologic symptoms of EAE, but SUW014 did not improve the symptoms. Data represent mean ± SEM; n=5 for vehicle and SUW014 (A) and n=9 and 11 for vehicle and TPPB (B), respectively. Statistical significance was determined by a two-tailed Student’s t-test for EAE clinical score; p-value of <0.05 was considered significant (* p<0.05 and ** p<0.01).

### TPPB Suppresses CNS Inflammation and Promotes Regeneration in EAE

To better understand the beneficial effect of TPPB on EAE, we examined the impact of TPPB on myeloid and lymphoid cell phenotypes in the CNS at peak EAE using flow cytometry. MOG_35-55_-induced EAE mice were treated with vehicle or TPPB (50 nmol/kg, three days per week) at the onset of clinical symptoms, and analyses were performed at peak disease on post-immunization day (PID) 19. Flow cytometry analyses revealed that treatment with TPPB suppressed CNS inflammation as demonstrated by a reduction in the population of CD11b^+^CD45^+^ cells expressing MHC class II in the spinal cord of EAE mice compared to the vehicle-treated group (Figure 3A). Furthermore, TPPB increased the population of CD11b^+^CD45^+^CD206^+^ cells in the spinal cord, suggesting that TPPB favors the activation of an anti-inflammatory/regenerative phenotype (Figure 3B). In addition, there was a significant downregulation in the proportion of CD4 lymphocytes displaying IL-17 (CD4^+^IL-17^+^) in the TPPB-treated group compared to the vehicle group, which indicates an efficient alleviation of inflammation by TPPB (Figure 3C). Overall, these results demonstrate that TPPB has a profound protective effect on the immune response in the MOG_35-55_-induced EAE model of MS.

**Figure 3.**
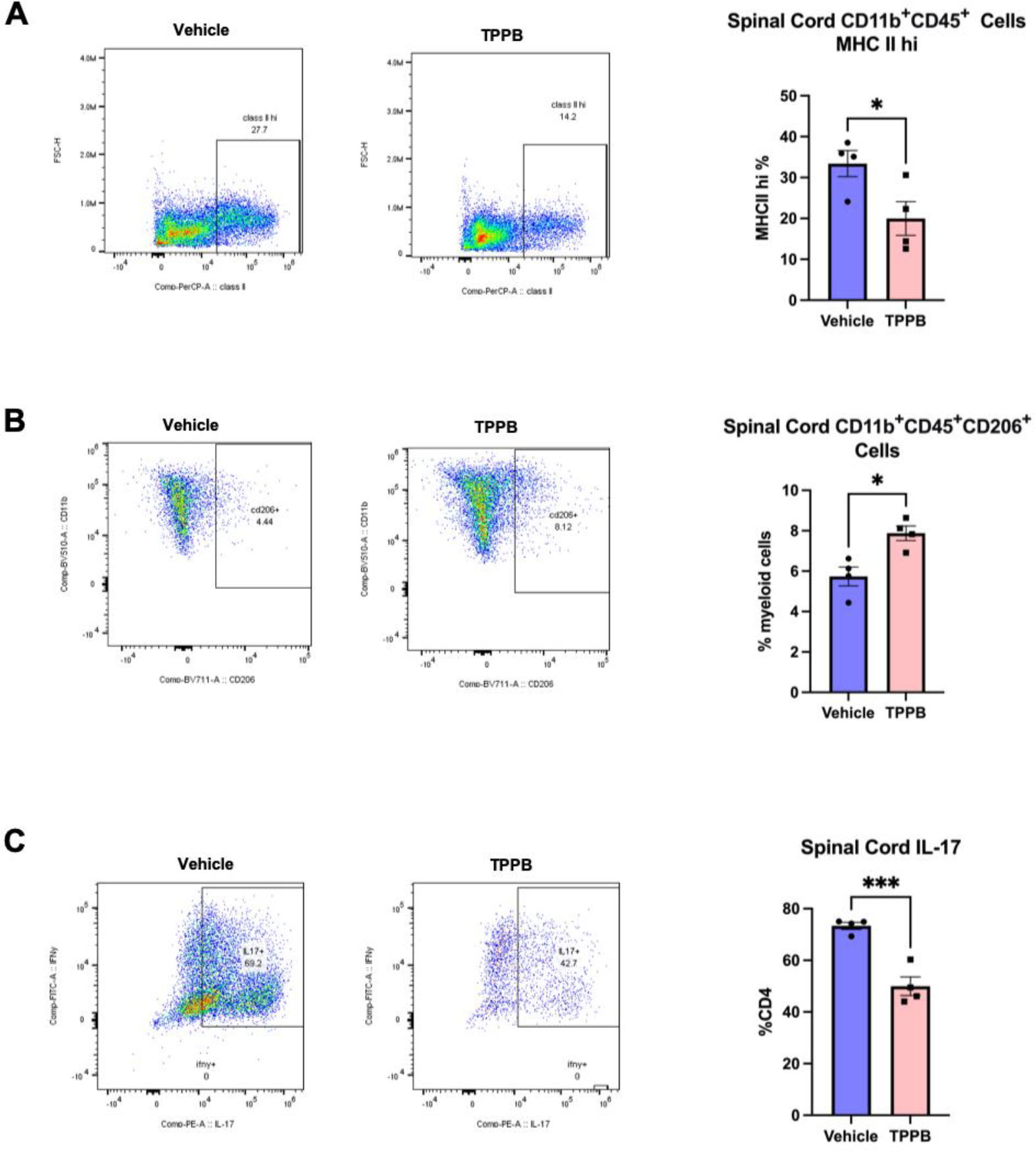
TPPB suppresses CNS inflammation in EAE mice. MOG_35-55_-induced EAE mice were treated with TBBP (50 nmol/kg, three days/week) or vehicle when neurologic symptoms first appeared, and they were sacrificed on PID 19. (A) Flow cytometry dot plots indicate the infiltrating CD11b^+^CD45^+^ cells examined for expression of MHC II and gated for MHC II hi cells in the spinal cords of the EAE mice (*left*). The bar graph (*right*) shows the decreased percentage of these cells in the TPPB-treated group as compared to the vehicle control group. (B) Flow cytometry dot plots show CD11b^+^CD45^+^CD206^+^ cells as percentage of CD11b^+^CD45^+^ cells in the spinal cord (*left*), and the bar graph (*right*) demonstrates the increase in the percentage of these cells in EAE mice treated with TPPB. (C) Flow cytometry dot plots show the expression of IL-17^+^ cells as percentage of total CD4^+^ T cells in the spinal cord of EAE animals (*left*), with the bar graph (*right*) indicating significant reduction in IL-17^+^ cells as a percentage of CD4^+^ T cells in the TPPB-treated group. n=4 for flow cytometry. All error bars represent SEM. Statistical significance was determined by a two-tailed Student’s t-test for flow cytometry data; p-value of <0.05 was considered significant (* p<0.05 and *** p<0.001).

### TPPB Potentiates the Repair Mechanism by Enhancing Phagocytosis

Having evaluated the potential of TPPB in the resolution of inflammation both in cultured BMDM and EAE animal model, we next sought to better understand the impact of TPPB on repair mechanisms by investigating its effect on phagocytosis. Phagocytosis is a key function of innate immune cells during remyelination by removing the deleterious and inhibitory myelin debris for an effective OL differentiation to occur. For the phagocytosis assay, BMDM were treated with either vehicle or TPPB (100 nM) for 24 hrs, and then were incubated with pHrodo E. coli BioParticles for 3 hrs followed by the measurement of relative fluorescence intensity on a plate reader. Compared to the vehicle condition, TPPB-treated group showed a significantly higher percentage of relative fluorescence intensity, suggesting increased phagocytosis when BMDM were treated with TPPB (Figure 4A). In addition, we performed a phagocytosis assay with CFSE-tagged myelin, where the BMDM were cultured on clear bottom 96-well plates and treated with either vehicle or TPPB (100 nM), followed by overnight incubation with CFSE-tagged myelin. ICW was performed to visualize the phagocytosed myelin, and β-actin staining was used to normalize the data (Figure 4B). Treatment of BMDM with TPPB significantly increased the phagocytosis of myelin (Figure 4C). The results from this experiment confirm that, in addition to the significant anti-inflammatory effect, TPPB also potentiates repair functions by promoting phagocytosis in innate immune cells.

**Figure 4:**
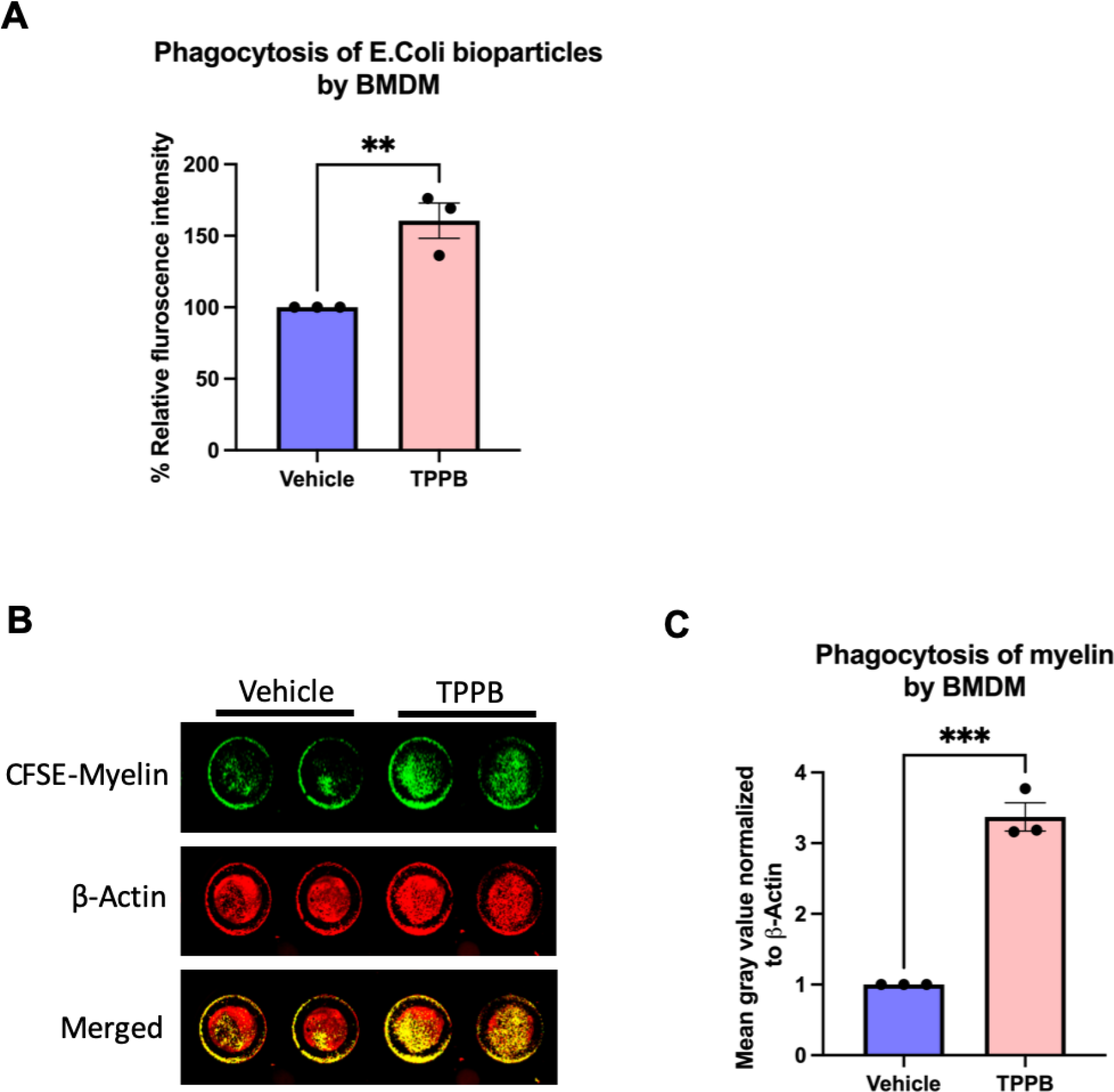
TPPB promotes phagocytosis in BMDM. (A) The bar graph representing the percentage relative fluorescence intensity shows significantly increased phagocytic activity of BMDM against E. coli particles when treated with TPPB (100 nM; n=3). All error bars depict SEM. (B) Representative image of ICW: BMDM were cultured in a 96-well plate and treated with either vehicle or TPPB (100 nM) followed by overnight incubation with CFSE-tagged myelin. β-actin was used as the loading control. (C) Calculated total fluorescence of ICW data quantified on ImageJ shows a significant increase in phagocytosis of CFSE-tagged myelin in the TPPB-treated group compared to the vehicle group. Quantification (mean ± SEM) from n=3 and statistical significance were determined by a two-tailed Student’s t-test; p-value of <0.05 was considered significant (** p<0.01 and *** p<0.001).

### TPPB Increases OL Differentiation Following Focal Demyelination

To evaluate the remyelination potential of TPPB in MS, we used the LPC-induced focal demyelination model. Focal demyelination was induced in the ventral spinal cord of the mice by stereotaxic injection of LPC. Treatment with TPPB (50 nmol/kg, three days/week) or vehicle was initiated 48 hrs post-LPC injection for two weeks. Representative images of lesions from 15 days post-lesion (dpl) in the TPPB- and vehicle-treated mice stained for Olig2^+^, CC1^+^, and DAPI are shown in Figure 5A. The number of total OL-lineage cells (Olig2^+^) did not change with TPPB treatment (Figure 5B). However, the number of the differentiating OL (Olig2^+^CC1^+^) within the lesions showed a trend towards increased differentiation in the TPPB-treated group when compared to the control group at 15 dpl (Figure 5C). The number of differentiating OL (Olig2^+^CC1^+^) represented as a percentage of total Olig2^+^ also showed a similar trend of increased OL differentiation in the TPPB-treated group (Figure 5D). This result suggests a potential enhancement of remyelination by TPPB, similar to bryo-1 (Gharibani *et al*., 2023).

**Figure 5:**
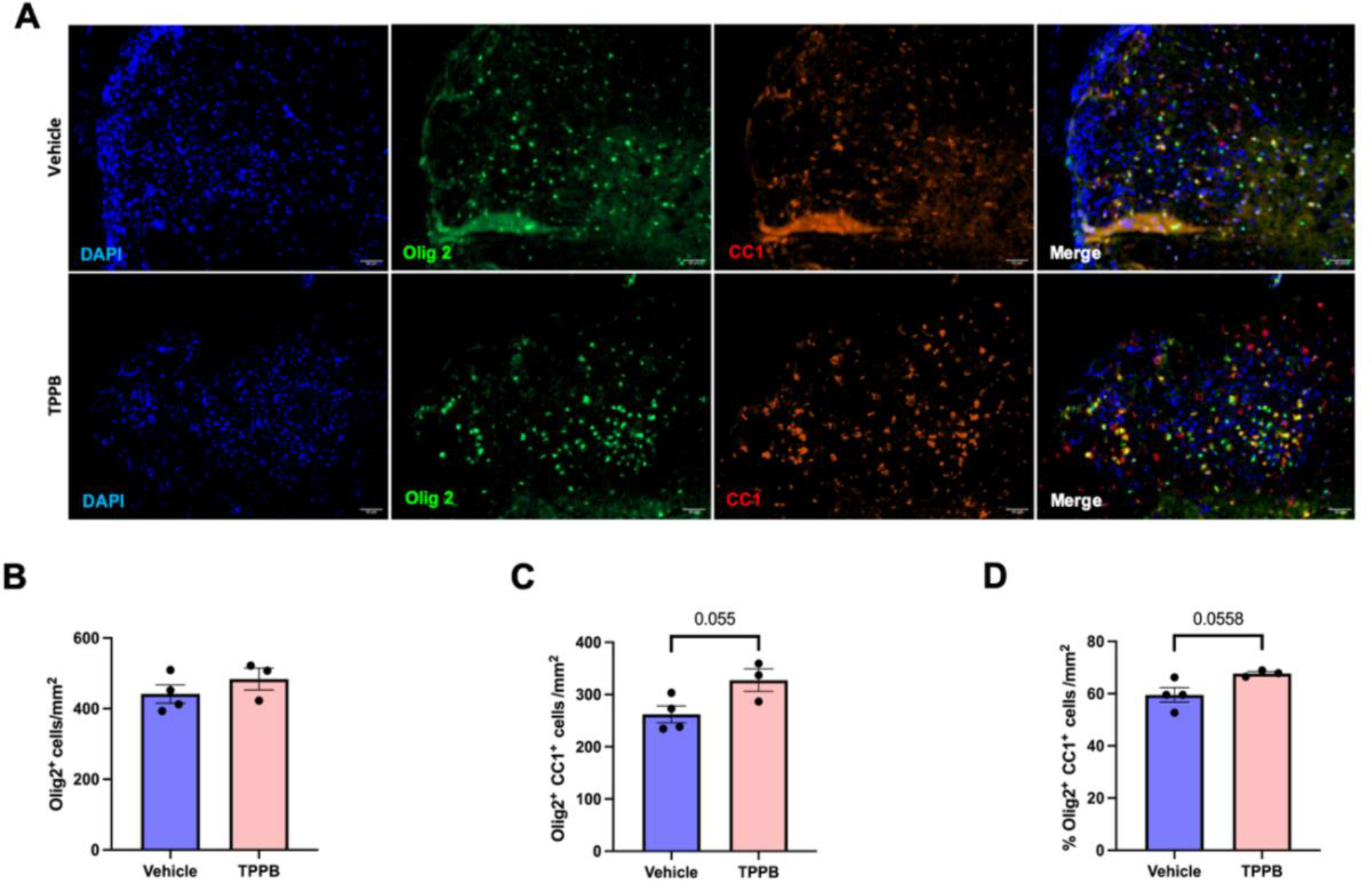
TPPB shows a trend towards enhancing OL differentiation following focal demyelination. (A) Representative images of LPC-induced demyelination lesions from 15 dpl in vehicle- and TPPB-treated mice stained for the shown markers. (B) Quantification of total OL-lineage cells (Olig2^+^) shows no significant difference in the total number of Olig2^+^ cells when treated with TPPB. (C) There is a trend towards increased differentiating OL (Olig2^+^CC1^+^) within the lesions at 15 dpl in TPPB-treated mice. (D) The bar graph shows the number of differentiating OL (Olig2^+^CC1^+^) represented as a percentage of Olig2^+^ cells, which also demonstrates similar trend as the total number of differentiating OL (B). Quantification (mean ± SEM) from n=4 mice for vehicle and n=3 mice for the TPPB group and statistical significance were determined by a two-tailed Student’s t-test; p-value of <0.05 was considered significant.

## Discussion

In our previous study, we demonstrated that our screening platform provided a rapid method for identifying PKC modulators for treating neuroinflammatory and neurodegenerative disorders (Abramson *et al*., 2021). That study showed that the prostratin analog SUW014 has a similar anti-inflammatory effect on cultured macrophages as bryo-1, and in this study, we have also identified TPPB as another potential PKC modulating compound with bryo-1-like properties. Our initial study primarily focused on bryo-1 and its analogs, but it did not further examine the in vivo effects of SUW014. Thus, to advance TPPB and/or SUW014 forward in our drug assay progression, we investigated the in vivo activity of these agents. We found that SUW014 interestingly did not ameliorate the neurologic symptoms of EAE, while TPPB was able to improve EAE phenotype like bryo-1 and bryologs. We recently established that bryo-1 is able to activate phagocytosis in macrophages and microglia (which is critical during remyelination as myelin debris inhibits myelin formation) and enhance remyelination in LPC-induced demyelination model (Gharibani *et al*., 2023). Here, we also demonstrated that TPPB stimulates phagocytosis by macrophages and increases remyelination in focal LPC-induced demyelination animal model.

The results from this current study indicate that while the in vitro immunologic assays are a useful initial screening method to identify candidate compounds, in vitro assays are not sufficient. As is found in many preclinical studies, in vitro activities do not always correlate with in vivo effects – in this case, a compound’s ability to mitigate neuroinflammation and promote remyelination. SUW014 demonstrates a greater potency than bryo-1 in triggering an anti-inflammatory response from in vitro macrophages, but it failed to alleviate EAE in vivo. While beyond the scope of this study but prompted by its findings, it would be important to determine why SUW014 was not able to treat the neurologic symptoms of EAE given that it had such promising in vitro activity. It is possible that SUW014 has poor bioavailability or was not absorbed adequately when administered IP. It is also feasible that SUW014 did not reach a therapeutic concentration in the CNS or that the dosing regimen of three times a week is insufficient. Loss through metabolism is another potential contributing factor as SUW014 has an esterase-labile ester functionality while TPPB has more stable amide bonds. Therefore, in vivo ester loss would eliminate PKC binding. It is also possible that off-target associations by SUW014 might contribute to the observed in vitro versus in vivo differences.

The discovery that TPPB and bryo-1 have similar in vitro and in vivo activities has several important implications. Our results demonstrate that TPPB, like bryo-1, has immunomodulatory effects in an EAE model, enhances phagocytosis, and increases remyelination. Identification of TBBP also indicates that the anti-inflammatory and regenerative properties of innate immune cells depend on PKC activity and not off-target effects of bryo-1, as it is unlikely that both bryo-1 and TPPB have the same off-target activities to provide these beneficial effects on innate immune cells. Additionally, our previous study established that modification of bryo-1 so that it can no longer bind to PKC (SUW275) prevents anti-inflammatory effects in myeloid cells and does not ameliorate EAE symptoms (Abramson *et al*., 2021). Finally, having structurally different chemicals with similar functional effects on PKC and in animal models of MS allows for diversification of candidates, which increases the probability of identifying the best candidate to advance forward in drug development for neuroinflammatory and neurodegenerative disorders.

PKC isoforms belongs to one of three classes, which are conventional (α, β, and γ), novel (δ, ε, η, and θ), and atypical (ζ and ι/λ). Conventional and novel PKCs have a C1 domain that binds to bryo-1, TPPB, and endogenous diacylglycerol (DAG), but atypical PKCs do not (Das and Rahman, 2014; Katti *et al*., 2022). Currently, it is unknown which isoform(s) is involved in anti-inflammatory and regenerative effects in microglia and CNS-associated macrophages. It is possible that TPPB intracellularly could activate more specifically the isoform(s) critical for immunomodulation of CNS innate immune cells, which would decrease the side effects that are typically observed with bryo-1, such as myalgia. The identification of PKC isoform(s) involved in this process and the specificity of TPPB and bryo-1 are under ongoing investigation.

Overall, we have discovered that TPPB, while significantly different in structure from bryo-1, has similar beneficial activities to bryo-1. This finding is consistent with their sharing common chemical properties that allow binding to PKC (Wender *et al*., 1986). TPPB serves as a new structural lead in an armamentarium of PKC-modulating drugs that could prove effective in the treatment of neuroinflammatory and neurodegenerative disorders for which there is a lack of therapeutic options. This study shows for the first time that the EAE activities of bryo-1 and its analogs are also exhibited in vivo by a structurally distinct molecular class, thus creating another therapeutic options for targeting CNS innate immune cells to modulate neuroinflammation and to promote CNS regeneration and repair.

## Author Contributions

Conceptualization: PMK

Data curation: SS, WHG, PG, PMK

Formal analysis: SS, WHG, PG, JJL, YG, XD, MDK, PMK

Funding acquisition: PAW, MDK, PMK

Investigation: SS, WHG, PG, JJL, PAW, MDK, PMK

Project administration: MDK, PMK

Resources: PAW, MDK, PMK

Supervision: MDK, PMK

Visualization: SS, WHG, PG, JJL, MDK, PMK

Writing – original draft: SS, PMK

Writing – review & editing: SS, WHG, PG, JJL, YG, XD, PAW, MDK, PMK

## Funding

Department of Defense, MS200232 (MDK, PMK)

TEDCO Maryland Innovative Initiative, 135025 (MDK, PMK)

NIH MSTP Grant T32 GM136577 (WHG)

NIH R01CA031845 (PAW)

American Association of Immunologists Careers in Immunology Fellowship (MDK, WHG)

## Acknowledgments

The authors thank L. Albacarys and P. A. Calabresi for their advice and support.

## Data availability Statement

All data are available in the main text or the supplementary materials.

